# Estimation of the dispersal distances of an aphid-borne virus in a patchy landscape

**DOI:** 10.1101/109561

**Authors:** David Pleydell, Samuel Soubeyrand, Sylvie Dallot, Gérard Labonne, Joël Chadœuf, Emmanuel Jacquot, Gaël Thébaud

## Abstract

Characterising the spatio-temporal dynamics of pathogens *in natura* is key to ensuring their efficient prevention and control. However, it is notoriously difficult to estimate dispersal parameters at scales that are relevant to real epidemics. Epidemiological surveys can provide informative data, but parameter estimation can be hampered when the timing of the epidemiological events is uncertain, and in the presence of interactions between disease spread, surveillance, and control. Further complications arise from imperfect detection of disease, and from the computationally intractable number of data on individual hosts arising from landscape-level surveys. Here, we present a Bayesian framework that overcomes these barriers by integrating over associated uncertainties in a model explicitly combining the processes of disease dispersal, surveillance and control. Using a novel computationally efficient approach to account for patch geometry, we demonstrate that disease dispersal distances can be estimated accurately in a fragmented landscape when disease control is ongoing. Applying this model to data for an aphid-borne virus (*Plum pox virus*) surveyed for 15 years over 600 orchards, we obtain the first estimate of the distribution of the flight distances of infectious aphids at the landscape scale. Most infectious aphids leaving a tree land beyond the bounds of a 1-ha orchard (50% of flights terminate within about 90 m). Moreover, long-distance flights are not rare (10% of flights exceed 1 km). By their impact on our quantitative understanding of winged aphids dispersal, these results can inform the design of management strategies for plant viruses, which are mainly aphid-borne.

**Author Summary:** In spatial epidemiology, dispersal kernels quantify how the probability of pathogen dissemination varies with distance. Spatial models of pathogen spread are sensitive to kernel parameters; yet these parameters have rarely been estimated using field data gathered at relevant scales. Robust estimation is rendered difficult by practical constraints limiting the number of surveyed individuals, and uncertainties concerning their disease status. Here, we present a framework that overcomes these barriers and permits inference for a between-patch transmission model. Extensive simulations show that dispersal kernels can be estimated from epidemiological surveillance data. When applied to such data collected from more than 600 orchards during 15 years of a plant virus epidemic our approach enables the estimation of the dispersal kernel of infectious winged aphids. This kernel is long-tailed, as 50% of the infectious aphids leaving a tree terminate their infectious flight within 90 m and 10% beyond 1 km. This first estimate of flight distances at the landscape scale for aphids–a group of vectors transmitting numerous viruses–is crucial for the science-based design of control strategies targeting plant virus epidemics.

## Introduction

Infectious diseases of humans, animals and plants severely impact the world’s health and economy. To gain knowledge on disease dynamics, powerful mathematical models have been developed [1–3]. However, for predicting the relative efficacies of competing control strategies across realistic heterogeneous landscapes, spatially-explicit *in silico* simulation models provide the main avenue [2]. The dispersal parameters of such models critically affect the predicted spatiotemporal dynamics of the disease, and thus the predicted outcome of potential control strategies [4]. Obtaining reliable estimates for these parameters is therefore a fundamental issue in epidemiology [5–7]. Models frequently employ dispersal kernels to represent how the probability of dispersal events diminishes as a function of distance [5]. Simulation studies have proven that dispersal parameters can be identified in idealised scenarios [5], which has been successful for simple models or small-scale datasets [8–13]. Recent advances in Bayesian methods and computing power have enabled fitting more realistic models to larger-scale surveillance data [6, 14–18]. However, most dispersal kernels are still often unknown. Indeed, estimation gets more complex when graduating from idealised toy problems to reconstructing the spatio-temporal dynamics of real epidemics. The first issue is the mismatch between the spatio-temporal coordinates of the epidemic, sampling and model [19]. For example, the timing of key events (e.g. when a susceptible individual becomes infected) is often censored (i.e. known only within certain bounds), and failure to account for this can bias estimates. Moreover, the challenge of inference is increased by uncertainty arising from missing observations [20, 21] or imperfect sensitivity of disease detection [22, 23]. Further difficulties arise when surveillance data are aggregated at the patch scale because a landscape comprising patches of various shapes or sizes often cannot be summarized by patch centroids without biasing connectivity estimates. All these issues require appropriate correction measures to avoid biased inference and prediction [24].

In the case of aerial vector- or wind-borne diseases, dispersal kernels critically depend on the flight properties of the vectors or infectious propagules [25]. When the probability of dispersal decreases more slowly than an exponential distribution, kernels are termed “long-tailed” and lead to non-negligible long-distance flights [26]. Such events are an important component of disease epidemiological–and evolutionary–dynamics and call for kernel estimation at the landscape scale [27]. However, among plant diseases, there are few available kernel estimates. The dispersal kernel of black Sigatoka (a fungal disease of banana) has been estimated experimentally up to 1 km of a point source, based on the direct observation of spore-induced lesions [28]. This is the only available direct estimate at this scale for the dispersal kernel of a plant disease, which reflects the extreme practical difficulties of such field studies and highlights the critical need for developing *in silico* solutions. A promising way forward is to infer dispersal parameters indirectly, i.e. from spatio-temporal patterns observed in epidemiological data [5] whilst accounting for the added complexity (outlined above) of observational studies. This approach has been used to infer the dispersal kernels of the wind-dispersed plantain fungus *Podosphaera plantaginis* [15], the fungus *Leptosphaeria maculans* affecting oilseed rape and dispersed both by wind and wind-driven rain [29], and two pathogens transmitted only by wind-driven rain: the oomycete *Phytophthora ramorum* that is responsible for sudden oak death [16], and the bacterium *Xanthomonas axonopodis* that causes Citrus canker [17]. A dispersal kernel has been estimated for two other Citrus diseases: Bahia bark scaling of Citrus, a disease with an elusive etiology [13], and Huanglongbing, which is caused by bacteria from the ‘*Candidatus* Liberibacter’ genus and transmitted by psyllids [18]. Up to now, this is the only vector-borne plant disease for which the dispersal kernel is documented. Although aphids are responsible for transmitting almost 40% of more than 700 plant viruses [30] and impose large economic burdens, their dispersal remains ill-characterized at the landscape scale [31, 32]. For a vast number of aphid-borne diseases, this lack of basic knowledge compromises the reliability of quantitative risk assessment and predictive epidemiological models–the key ingredients of science-based control strategies.

Most aphid-borne viruses belong to the *Potyvirus* genus and are transmitted in a non-persistent manner, i.e. by winged aphids that acquire and transmit the virus immediately while probing on various plants in search of a suitable host species [30]. Potyviruses are transmitted by a wide range of aphid species, and aphid infectivity is lost after the first probes. For these reasons, estimating the natural dispersal kernel of a potyvirus provides an indirect way of estimating the dispersal kernel of infectious winged aphids. *Plum pox virus* (PPV) is a potyvirus that appears in a list of the 10 most important plant viruses [33]. This virus is the causal agent of sharka, a quarantine disease affecting trees of the *Prunus* genus (i.e. mainly peach, apricot and plum), reducing fruit yield, quality (modified sugar content and texture) and visual appeal (due to deformations and discolouration) [32]. Sharka is a worldwide plague that has infected over 50 countries in Europe, Asia, America and Africa [32], inflicting estimated economic losses of 10 billion Euros over 30 years [34]. The transfer of infected (possibly symptomless) plant material can disseminate PPV over long distances [34], and the natural spread of the disease is ensured by more than 20 aphid species [35]. Virus-infected trees cannot be cured, and insecticides do not act fast enough to prevent the spread of the virus by non-colonising aphids [30, 36]. In addition, resistant or tolerant peach and apricot varieties are too scarce to provide a short-term alternative to the cultivated varieties. However, local aphid transmission can be reduced by removing infected trees as soon as they are detected. As a result, various countries have implemented PPV eradication or control strategies based on regular surveys and removal of trees or orchards when PPV is detected [32,34,37]. Given the cost of surveillance, tree removal and compensation, these strategies should benefit from model-assisted optimisation, which requires estimating the aphid dispersal kernel.

In this context, the aims of this study are: (i) to develop a Bayesian inference framework for estimating from surveillance data the parameters of a spatially-explicit epidemiological model that accounts for patch geometry and for interactions between disease spread, surveillance and control, (ii) to assess through simulations the accuracy and precision of the dispersal parameters estimated under various epidemic scenarios, and (iii) to apply our method to 15 years of geo-referenced surveillance data collected during an epidemic of *Plum pox virus* in order to estimate the dispersal kernel of the aphid vectors.

## Materials and Methods

### Surveillance database

In the early 1990’s, an outbreak of the M strain of PPV was detected in peach/nectarine patches (orchards) in southern France [38]. The plant health services implemented a control strategy based on disease surveillance and removal of symptomatic trees. This process involved the routine collection of patch-level data comprising the observed number of new cases (trees with PPV-typical discolouration symptoms on flowers and leaves) and the corresponding inspection dates, as well as patch attributes (location, planting and removal years, planting density, etc.). We aggregated the information for surveillance years 1992-2006 into a unique georeferenced database, with patch boundary coordinates obtained from digitised aerial photographs. With 4820 inspections over 15 years in 605 patches (52 of which were replanted during that period), this database is a precious resource for inference on aphid-mediated viral dispersal in fragmented landscapes. Moreover, to account for seasonal variations in the number of flying aphids, we used in our model the average (over 17 years) weekly number of flying aphids collected from a 12-m-high Agraphid suction tower located within the bio-geographical region of the study area.

### Modelling Framework

Our model has a compartmental Susceptible-Exposed-Infectious-Removed (*SEIR*) structure that aims to reduce bias in parameter estimates by accounting for irregular patch geometry, detection-dependent removal, imperfect detection sensitivity, interval censoring of between-compartment transition times, missing data and parameter uncertainty. We address these challenges by: (i) integrating a mixture of exponential dispersal kernels over source and receiver patches to compute between-patch connectivity; (ii) splitting the infectious state *I* into hidden (*H*) and detected (*D*) sub-states (Figure 1); (iii) integrating over uncertainty in the times of transition between compartments; (iv) using Bayesian data augmentation and inference. Two versions of our discrete-time spatio-temporal *SEHDR* model (one for stochastic simulations and the other for Bayesian inference) are described below (for further details, see Texts S1 and S2).

**Figure 1.**
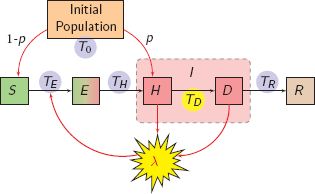
Susceptible-Exposed-Hidden-Detected-Removed (*SEHDR*) model of an individual’s epidemiological status. At *T*_0_, patch *i* is planted with infectious (*I*) or susceptible (*S*) individuals with probabilities *p*_*i*_ and 1-*p*_*i*_, respectively. An individual passes between compartments at event times *T*_*E*_, *T*_*H*_, *T*_*D*_ and *T*_*R*_. Only the detection time *T*_*D*_ is known; all other event times are censored. Infectious individuals from both within and outside the patch contribute to the force of infection *λ*_*t*_*r*, which is the expected number of infectious events affecting an individual over time interval (*t*_*r-*1_*, t*_*r*_]. The probability that a given susceptible (*S*) individual becomes exposed (*E*) in this time interval is 1-exp(-*λ*_*t*_*r*), assuming independent infection events. A latent period of duration *T*_*H*_ -*T*_*E*_ follows, after which the individual becomes infectious (*H*). Infectious individuals are removed (*R*) only after detection (*D*) or when the entire patch is removed. For simplicity, the *i* and *t*_*r*_ subscripts are omitted in the figure.

### Model Structure

Each patch *i* is planted with *N*_*i*_ individuals. At the planting date, a proportion *p*_*i*_ of these individuals are infectious (in state *H*) and 1-*p*_*i*_ are susceptible (in state *S*). If patch *i* is an introduction patch, *p*_*i*_*>*0; otherwise, *p*_*i*_=0. Up to four transition times (*T*_*E*_*, T*_*H*_ *, T*_*D*_ and *T*_*R*_) can be associated with any given individual (Figure 1), i.e. individuals pass sequentially from state *S* to *E* to *H* to *D* to *R*, and all other transitions occur with zero probability. The exposed state *E* accounts for the latent period, i.e. the time lag between the infection date *T*_*E*_ and the date at which the individual becomes infectious *T*_*H*_. In this discrete-time model (whose time steps are denoted by the index *r*), the transitions (denoted by ‘→’) between the five compartments are modelled as:

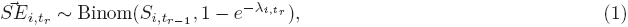

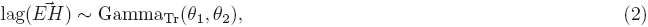

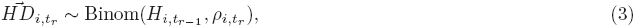

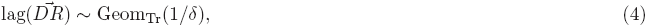

where: *S*_*i,tr−*1_ (resp. *H*_*i,tr−*1_) is the number of individuals in patch *i* that are in state *S* (resp. *H*) at the beginning of the time interval (*t*_*r*-1_*, t*_*r*_], and 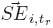 represents how many of them make the transition from *S* to *E* (resp. from *H* to *D*) in this time interval; the sojourn times in compartments *E* and *D* are determined per individual via random variables 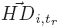 and 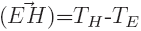, respectively; *ρ*_*i,*_*t*_*r*_ is the probability of detecting symptoms on an infectious (*H*) individual (*ρ*_*i,t*_r =*ρ* when patch *i* is inspected in (*t*_*r*-1_*, t*_*r*_], and 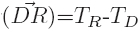 otherwise); the left truncation of Gamma_Tr_ represents an absolute minimal latent period for sharka [32]; the right truncation of Geom_Tr_ (with mean *δ*) represents the usual maximal delay between detection and removal. The force of infection (i.e. the expected number of transmission events) incurred by each individual in patch *i* over (*t*_*r*-1_*, t*_*r*_] is defined as:

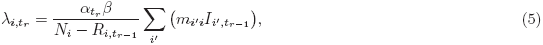

where *α_t_r__* is the normalized flight density, i.e. the proportion of annual flights occurring over (*t*_*r*-1_*, t*_*r*_]; *β* is the transmission coefficient, i.e. the annual number of vector flights per source (infectious) host that would lead to infection if the recipient host is susceptible; *N*_*i*_−*R*_*i,tr−*1_ is the number of remaining hosts in patch *i* and *I*_*i* ′, *tr* − 1_ is the number of infectious hosts in patch *i*′ over (*t*_*r*-1_*, t*_*r*_]. Note that *N*_*i*_ is constant (i.e. *N*_*i*_ = *S*_*i,tr*_ + *E*_*i,tr*_ + *I*_*i,tr*_ + *R*_*i,tr*_) for all *t*_*r*_ between the planting and removal dates of patch *i*. Finally, the connectivity *m*_*i*_′ *i* is the probability that a vector flight starting in patch *i*′ terminates in patch *i*.

The mean connectivity between source patch *i*′ of area *A*_*i*_′ and receiver patch *i* [39] is obtained via:

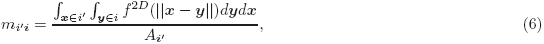

where **x** and **y** are coordinate vectors in ℝ^2^, and *f* ^2*D*^ is the 2-dimensional dispersal kernel. The computation time required to calculate connectivity *m*_*i*_′ *i* between several hundreds of patches prohibits the use of iterative algorithms to directly estimate the parameters of flexible (e.g. exponential-powered) kernels. Thus, we developed an approach that can approximate many long-range dispersal kernels. We defined *f* ^2*D*^ as a mixture of *J* components:

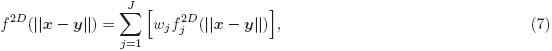

where the *w*_*j*_ are positive mixture weights summing to 1, and 2*h*_*j*_ is the mean dispersal distance for exponential kernel 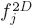defined as:

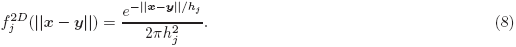

Under this mixture formulation, the connectivity becomes:

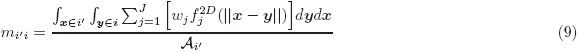

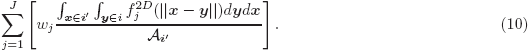

This permits the connectivity of each mixture component *j* to be computed just once, and only the weights *w*_*j*_ need to enter the iterative estimation procedure. We set 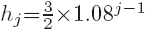(and *J*=100), to obtain kernel components with mean distances ranging from 3 to 6110 m, and higher resolution at smaller distances. To simplify parametrisation, and to avoid identifiability issues with the mixture of exponentials, we restrain weights using:

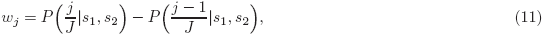

where *P* is the cumulative density function of a beta distribution with parameters *s*_1_ and *s*_2_. We call any kernel of the form (eq. 7) weighted by eq. 11 a beta-weighted mixture of exponentials (BWME) kernel.

### Estimation Model

We now focus on the model used for parameter estimation. Among the four transition times, only *T*_*D*_ (i.e. the time when an infectious individual is detected) is known precisely. Let *(t*_*i*,1_,⋯, *t*_*i,k,*_⋯, *t*_*i,Ki*_) denote the set of *K*_*i*_ inspection dates in patch *i* (which may be partly censored by omissions in the surveillance records). Let *p*(*T*_*D,i*_ = *t*_*i,k*_) denote the probability for an individuals in patch *i* to be detected as infected at inspection date *t*_*i,k*_. Data provide the associated number 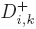 of newly detected individuals, which is modelled as:

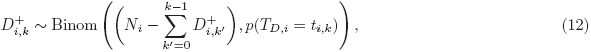

with 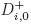 = 0. To account for censoring and imperfect detection sensitivity, a survival model was used to derive *p*(*T*_*D,i*_ = *t*_*i,k*_) (Text S1). Using eq. (12) we obtained the likelihood of the observed data given a set of parameters Θ (Text S1). Based on this likelihood, Bayesian inference of parameter distributions was performed via a Gibbs sampler with embedded adaptive Metropolis-Hastings steps (Text S3).

### Estimation for Simulated Epidemics

To assess the accuracy (i.e. lack of bias) and precision (i.e. lack of variance) of the estimation of the dispersal parameters, 10 epidemics were simulated under each combination of 7 disease introduction scenarios × 3 dispersal kernels × 4 parameter estimation scenarios. All simulations were performed under the same virtual landscape derived from the surveillance database: we retained the spatial coordinates (and thus the area) of the patches, but the other potential spatio-temporal dependencies were suppressed through the random permutation of planting densities and of patch planting/removal/replanting dates. When density or planting date were missing in the database, their values were drawn from the corresponding empirical distribution. Simulations were performed with 1 time step per day, and 1 survey per patch per year, with inspection days drawn from the corresponding empirical distribution. The transmission coefficient *β* was fixed at 1.5 (which leads to realistic epidemic dynamics) and all other parameters were fixed at the expected values of their prior distributions (Text S2).

The three simulated kernels correspond to short-, medium-and long-range dispersal. They were parametrised using low-dimension mixtures of exponential kernels (eq. 7) with fixed mean distances and weights (Table 1, mixture parameters). These were subsequently approximated by the BWME kernel minimizing the Kullback-Leibler (KL) distance between the two probability density functions (Table 1, simulation parameters).

**Table 1.**
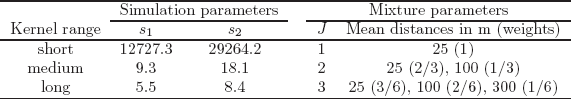
Parameters for 3 simulated dispersal kernels. Epidemics were simulated using shape parameters *s*_1_ and *s*_2_ for the BWME kernel (left) approximating simpler exponential mixture kernels with just *J* mixture components (right).

The seven introduction scenarios were defined by the following number of introduction patches (and the initial prevalence *p*_*i*_ in these patches): 1 (25%), 5 (10%), 10 (5%), 15 (2%), 20 (1%), 25 (1%) or 30 (1%). For a given introduction scenario, all simulations were performed with the same introduction patches, which were chosen at random with the constraint that the first introduction occurred at year 1 and all other introductions occurred before year 6 (Figure S1).

In order to identify whether our Bayesian Markov chain Monte Carlo (MCMC) estimation procedure (Text S3) encountered identifiability issues with some parameters, we tested 4 estimation scenarios targeting parameter sets of increasing size (Table 2), with all other parameters fixed at the values used for simulation.

**Table 2.** Parameter sets for the 4 estimation scenarios. For each estimation scenario, the parameter set Θ comprises the parameters indicated with a **✓**.

| Parameter | Definition | $\Theta_1$ | $\Theta_2$ | $\Theta_3$ | $\Theta_4$ |
| --- | --- | --- | --- | --- | --- |
| $\beta$ | transmission coefficient | ✓ | ✓ | ✓ | ✓ |
| $\mu$ | $=s_1/(s_1+s_2)$; mean of kernel weight distribution | ✓ | ✓ | ✓ | ✓ |
| $\sigma$ | $=s_1+s_2$; shape of kernel weight distribution | ✓ | ✓ | ✓ | ✓ |
| $\rho$ | detection sensitivity | - | ✓ | - | ✓ |
| $\theta_1$ | shape of latent period distribution | - | - | ✓ | ✓ |
| $\theta_2$ | scale of latent period distribution | - | - | ✓ | ✓ |

Both simulation and estimation algorithms started at the beginning of year 1 and stopped at the end of year 22. Running 10 MCMC chains under each estimation scenario applied to each simulated epidemic produced 8400 MCMC chains in total. Within each epidemic replicate × kernel × introduction × estimation scenario combination, we retained the MCMC chain with the highest mean posterior log-likelihood. Then, for each of these 840 best chains, indices of accuracy (resp. precision) were defined as the mean (resp. span of the 95% credibility interval) of the posterior KL distances between the probability density functions *f* ^2*D*^ (eq. 7) of simulated and estimated kernels. Simulated and estimated kernels were plotted using the distribution function of the distance travelled:

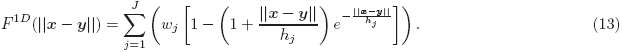

This function is the cumulative version of the 1-dimensional *f* ^1*D*^ (i.e. the probability density function of the distance travelled), which was obtained by integrating (marginalising) *f* ^2*D*^ (eq. 7) over all directions.

### Estimation for a Real Epidemic

Using PPV surveillance data, we applied our estimation algorithm to infer parameters of the spatial *SEHDR* model from the MCMC chain (among 10) with the highest mean posterior log-likelihood (Text S3). The number of introduction patches *κ* was fixed at integer values in the range 1-25, and 30 chains were run per fixed *κ*. This approach was taken because each unit increase in *κ* adds two parameters (additional introduction patch identity and initial prevalence) to Θ, which always increases the posterior log-likelihood (various uninformative and weakly informative priors were tested). Thus, to avoid over-fitting, identification of *κ* was treated as a model selection problem for which we maximised the Fisher information criterion I(*κ*) (Text S4).

## Results

### Impact of Parameter Values on Simulated Epidemics

The parameter combinations chosen to test the inference procedure cover a wide range of epidemic behaviour, from local to widespread epidemics and from low to high incidence (Figure 2). The general trends are that more introduction patches unsurprisingly lead to more widespread epidemics, and that higher disease prevalence in the introduction patches does not increase much the final local prevalence. As expected, increasing kernel ranges generally causes more widespread epidemics but with decreasing prevalence in the infected patches (in particular, close to the introduction patches).

**Figure 2.**
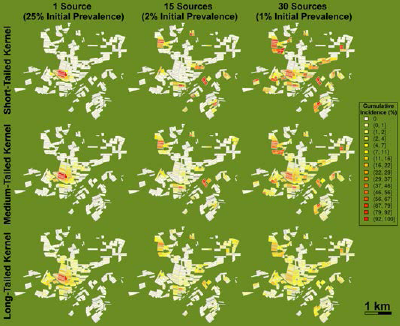
Cumulative detected incidence at the end of year 22 for nine typical simulated epidemics. Each polygon represents one peach orchard. From left to right, the number of introduction patches (with initial disease prevalence) are: 1 (25%), 15 (2%) and 30 (1%). From top to bottom: simulations generated under short-, medium-and long-range kernel scenarios.

### Evaluation of the Estimation Procedure

The distribution of KL distances between simulated and estimated kernels demonstrates that estimation accuracy is not affected by the inclusion of sensitivity and latent period parameters in the estimation scheme (Figure 3A). The median accuracy of the estimated kernels is not much affected either by the range of the dispersal kernel (Figure 3B). However, for longer-range dispersal kernels, KL distances can become more extreme (Figure 3B), and the span and variance of their 95% credibility intervals increase (Figure S2B). This shows that the precision of the estimated kernel decreases with increasing dispersal range. The most influential factor on accuracy and precision of estimated dispersal kernels is the introduction scenario (panel C in Figures 3 and S2). However, the effect of the introduction scenario is not strongly related to the number of introduction patches or the associated initial prevalence, but rather to the presence of an introduction patch in the dense central cluster of patches (Figures 3 and S1).

**Figure 3.**
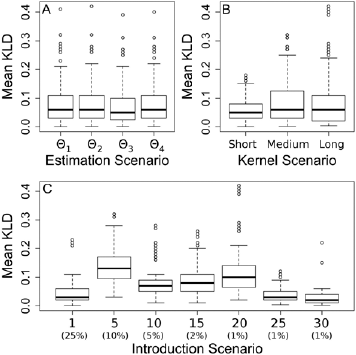
Boxplot of the distances between simulated and estimated dispersal kernels. Impact of (A) estimation scenario, (B) kernel range, and (C) disease introduction scenario [number of introduction patches (with initial disease prevalence)] on the accuracy of estimated dispersal kernels. Accuracy is measured by the Kullback-Leibler distance between simulated and estimated dispersal kernels.

For each of the 3 simulated kernels, the distribution of KL distances was summarised by its minimum, quartile and maximum values across all 7 introduction scenarios × 10 epidemics per scenario. The comparison between simulated kernels and their estimates within the richest scheme (Θ_4_) shows that the 3 kernels are very accurately estimated for some simulated epidemics (Figure 4, left column). However, dispersal distances are often overestimated, with the median KL distance increasing from 5.2×10^-2^ to 6.1×10^-2^ with increasing kernel range. A closer look at the median estimation curves reveals that the estimated distances never exceed the simulated distances by more than 0.25 on the logarithmic scale. Dispersal distances are thus overestimated by a factor below 1.8. Even for the most challenging of the 70 epidemics simulated with the long-range dispersal kernel (bottom-right panel), the difference between the two curves remains below 0.6 on the logarithmic scale. This value translates into less than 4-fold estimation errors, which is high but still within one order of magnitude. By contrast, precision is very high for all kernel ranges, as indicated by a median span below 0.04 for the 95% posterior credibility interval of KL distances (Figure S2) and the corresponding overlapping red lines in each plot of Figure 4. The results are similar for the other estimation scenarios (not shown).

**Figure 4.**
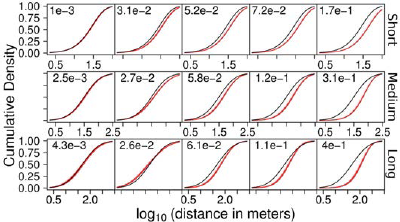
Comparison of simulated and estimated dispersal kernels. From left to right: kernels with the minimum, lower quartile, median, upper quartile and maximum Kullback-Leibler (KL) distances (posterior mean), as estimated (red) under the richest parameterisation scheme (Θ_4_), based on simulated epidemics with short-, medium-and long-range kernels (from top to bottom; black). Kernels are represented by their marginal cumulative distribution function *F* ^1*D*^ (with distance from the source represented on the log_10_ scale). The mean KL distance is indicated for each estimation.

### Estimation for a Real Epidemic

Once validated on simulated epidemics, we used the developed inference framework to estimate the dispersal kernel of *Plum pox virus* (and thus of the flight distances of the infectious aphid vectors) based on survey data. As a first step, we inferred the number of introduction patches. For *κ<*10, no combination of introduction patches returned a finite posterior log-likelihood. The Fisher information criterion was maximised at *κ*=11 (Figure 5), indicating that improvement in model fit saturates beyond this point. This suggests that the most robust inference is obtained with *κ*=11. These 11 introduction events among 547-579 orchards planted over 22 years (planting date is unknown for 32 orchards) correspond to disease introduction probabilities of 0.5 per year and 1.90-2.01×10^-2^ per orchard planted.

**Figure 5.**
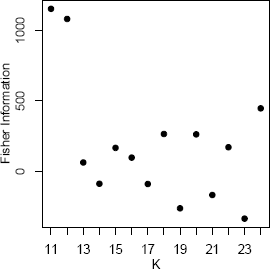
Impact of the number of introduction patches (*κ*) on the expected Fisher information for the sharka epidemic. For each *κ*, the estimation with the highest mean posterior log-likelihood was retained. For *κ<*10 no introduction patch combination returned a finite posterior log-likelihood. The empirical approximation of the Fisher information was maximal at *κ*=11.

Summary statistics of the posterior distributions of key parameters and percentiles of the dispersal kernel were tabulated for *κ*=11 (Table 3). From the estimated values of *s*_1_ and *s*_2_, we derived the weights of the kernel components (Figure S3), the dispersal kernel and the cumulative distribution function of aphid flight distances (Figure 6). This figure, and the estimated quantiles shown in the second part of Table 3, demonstrate the substantial contribution of long-range dispersal to aphid-borne virus epidemics. Indeed, almost 50% of the infectious aphids leaving a tree land beyond 100 m (median distance = 92.8 m; *CI*_95%_=[82.6-104 m]), and nearly 10% land beyond 1 km (last decile = 998 m; *CI*_95%_=[913-1084 m]).

**Table 3.**
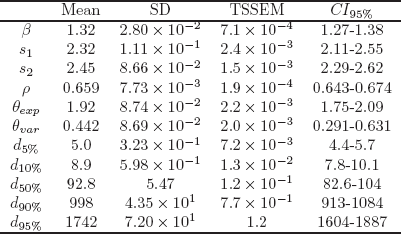
Summary statistics of marginal posterior distributions for the real epidemic. Summary statistics including the mean, standard deviation (SD), time-series standard error of the mean (TSSEM) and 95% credibility intervals (CI_95%_) are reported for the transmission coefficient *β*, the kernel parameters *s*_1_ and *s*_2_, the detection sensitivity *ρ*, the expected duration *Θ*_*exp*_ of the latent period and associated variance *Θ*_*var*_. Posterior distributions of the 5^th^, 10^th^, 50^th^, 90^th^ and 95^th^ percentiles of aphid flight distances *d* are also summarised.

| | Mean | SD | TSSEM | $CI_{95\%}$ |
| --- | --- | --- | --- | --- |
| $\beta$ | 1.32 | $2.80 \times 10^{-2}$ | $7.1 \times 10^{-4}$ | 1.27-1.38 |
| $s_1$ | 2.32 | $1.11 \times 10^{-1}$ | $2.4 \times 10^{-3}$ | 2.11-2.55 |
| $s_2$ | 2.45 | $8.66 \times 10^{-2}$ | $1.5 \times 10^{-3}$ | 2.29-2.62 |
| $\rho$ | 0.659 | $7.73 \times 10^{-3}$ | $1.9 \times 10^{-4}$ | 0.643-0.674 |
| $\theta_{exp}$ | 1.92 | $8.74 \times 10^{-2}$ | $2.2 \times 10^{-3}$ | 1.75-2.09 |
| $\theta_{var}$ | 0.442 | $8.69 \times 10^{-2}$ | $2.0 \times 10^{-3}$ | 0.291-0.631 |
| $d_{5\%}$ | 5.0 | $3.23 \times 10^{-1}$ | $7.2 \times 10^{-3}$ | 4.4-5.7 |
| $d_{10\%}$ | 8.9 | $5.98 \times 10^{-1}$ | $1.3 \times 10^{-2}$ | 7.8-10.1 |
| $d_{50\%}$ | 92.8 | 5.47 | $1.2 \times 10^{-1}$ | 82.6-104 |
| $d_{90\%}$ | 998 | $4.35 \times 10^1$ | $7.7 \times 10^{-1}$ | 913-1084 |
| $d_{95\%}$ | 1742 | $7.20 \times 10^1$ | 1.2 | 1604-1887 |

**Figure 6.**
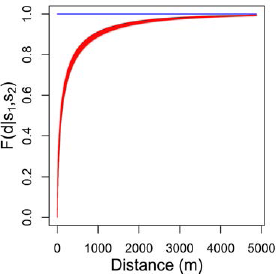
Estimated dispersal kernel for the sharka epidemic. Posterior distributions of dispersal kernels are represented by their marginal cumulative distribution function *F* ^1*D*^ obtained for *κ*=11 (i.e. the number of introduction patches maximising the Fisher information). The plotted posteriors were obtained from 4000 samples of the MCMC chain with the highest mean posterior log-likelihood.

## Discussion

In this work, we have developed a spatially-explicit Bayesian inference framework for the estimation of disease dispersal parameters when surveillance data are gathered at the patch level. The simulation and inference procedures take into account that disease status assessment is incomplete because surveillance has an imperfect detection sensitivity and a finite spatio-temporal coverage. We assessed the quality of the inference procedure through the comparison between parameter values used for simulation and the corresponding estimates. Then, we applied this approach to *Plum pox virus* surveillance data, to obtain the first estimate of an aphid dispersal kernel at the landscape scale. We discuss below the interest and limitations of the proposed approach and results.

### Sources of Uncertainty and Model Validation

We have focused attention on the estimation of the dispersal kernel since this is the key component of spatial epidemiological models. Recent methodological advances have permitted to extract from surveillance data crucial information on the dispersal kernel of four plant diseases [13,16–18]. These estimation procedures all account for unobserved infection times, with additional methodological challenges related to large heterogeneous landscapes [16], introduction from external sources [17, 18], or active disease control [18]. The present work handles these various processes and, contrary to the abovementioned studies which all assume a known detection sensitivity, also accounts for this unknown variable which adds a layer of uncertainty into the surveillance process. Inclusion of parameters for detection sensitivity and the latent period in the estimation procedure (Table 2) barely affects the KL distance between simulated and estimated kernels (Figures 3 and S2). Hence the inclusion of these extra parameters during parameter inference based on PPV surveillance data.

A unique feature of the present work is the validation of the estimation of the dispersal kernel through comparing known functions used in simulations and the corresponding functions estimated from these simulated epidemic data. Although this is an intuitive and standard practice, previous studies instead used goodness-of-fit statistics between actual and simulated spatiotemporal patterns as a way to validate their models [16–18]. One reason for this discrepancy is that we specifically focus on the estimation of the dispersal kernel rather than on model predictions (as in [16, 17]). Another likely reason is the high computational burden associated with simulation-based validation procedures which require testing estimation algorithms under several simulation scenarios, with several independent estimations per scenario, to assess robustness, accuracy and precision. Here, this procedure proved useful to demonstrate that dispersal kernel estimation is generally more accurate for shorter-range kernels and depends on the epidemic dynamics, leading either to very precise estimation, or to overestimation, of dispersal distances (Figures 3 and 4).

### Dispersal Estimation at the Landscape Scale

Our inference procedure explicitly accounts for patch geometry and patch-level aggregation of surveillance data. Although this choice was data-driven (infected tree numbers–not locations– were included in the database), for landscape-scale studies this approach appears to strike an interesting compromise between computational feasibility and spatial realism. Indeed, although disease dynamics over the full landscape is informative on long-range dispersal, considering disease status of over 300,000 individuals simultaneously would cause major computational issues. Conversely, summarizing patch layout by patch centroid coordinates (as often done in spatial models) can bias connectivity estimates, especially when patch shapes are disparate, and when distances between patches are of the same order of magnitude as patch dimensions. The proposed approach can thus be useful to estimate the lanscape-scale dispersal kernels of many wind-and vector-borne diseases.

A rigourous assessment of connectivity between patches is also necessary because of its influence on parameter estimation. Indeed, our study shows that the KL distance between simulated and estimated dispersal kernels is affected by kernel range (Figures 3 and S2). This pattern reflects how parameter identifiability depends on statistical power, which depends on cumulative disease incidence, which in turn depends on landscape connectivity. Short-range kernels imply greater local connectivity than long-range kernels, leading to relatively intense local transmission but reduced transmission at greater distances. Whether or not shorter-range kernels generate larger epidemics depends on the proportion of potential transmission events falling outside host patches, and thus on landscape configuration. Here, larger cumulative incidences were obtained using smaller kernels because we worked with a fragmented agricultural landscape.

### Impact of Disease Introductions on Inference

Disease introduction scenarios had a substantial effect on the accuracy and precision of the inferred dispersal kernel (Figures 3 and S2). Surprisingly, this effect does not seem related to either the number of introduction patches or the associated initial prevalence. However, we note that lower KL distances between simulated and estimated dispersal kernels (in introduction scenarios 1, 6 and 7) are associated with introductions occurring in the highly connected central patches (Figure S1). The resulting higher cumulative incidence probably improves estimation for the reasons given above.

During parameter estimation, we did encounter multi-modality in the posterior likelihoods, which often happens when observed patterns are only indirectly related to the modelled processes. For epidemic scenarios with both a short-range kernel and a high number of introduction events, misidentifying some of the introduction patches had a large negative effect on the likelihood. For this reason, we ran the MCMC algorithms many times and carefully compared the posterior likelihoods and parameter estimates of all chains before making inference. We also considered alternative algorithms such as parallel tempering [40] or equi-energy sampling [41], but the extra computational burden of these approaches was considered superfluous given that the observed differences in the posterior likelihoods of various modes were typically relatively large. Thus, launching a large number of chains clearly increased the likelihood of identifying the global mode. We have extensively tested this approach, reporting here the results of 8880 chains, and have found that in practice results are consistent.

Overall, inference of epidemiological parameters is easier for epidemics where disease introductions are well characterized, or at least infrequent. Unfortunately, this was not the case with the PPV-M dataset, and estimating the number of introduction patches *κ* was challenging. Such difficulty is by no means unique to the current study (see e.g. [17]). Reversible-jump MCMC (RJMCMC) [42] is a popular solution to a similar problem arising in mixture models. We initially attempted various implementations of RJMCMC, but found it impossible to construct priors that could both prevent over-fitting and provide robust posterior probabilities for *κ* under a wide variety of epidemiological scenarios. To circumvent this issue we inferred *κ* based on the Fisher information. This gives a minimum-variance estimator that provides robust inference with a good balance between under-and over-fitting –although it does not permit the estimation of posterior probabilities associated with the various *κ*. This approach has been used successfully in similar situations [43].

### Insights into Aphid Biology

Like most plant viruses, PPV is transmitted in a non-persistent manner by winged non-colonising aphids [32]. The distance travelled by an aphid within a single flight is thus crucial to plant virus epidemiology. However, this dispersal kernel has long remained elusive. Traditional ecological methods such as capture-mark-recapture provide little information regarding aphid dispersal at the landscape scale [31]. This has been a major obstacle to the parametrisation of models simulating the dispersal of these vectors and the pathogens they spread, as exemplified by the scarcity of landscape-scale models on cereal aphids [44] and by the informed guesses of flight-distance parameters in such models [45]. Here we estimated, for the first time, the dispersal of aphid vectors at the landscape scale. This estimation indicates that 50% of the infectious aphids leaving a tree land within about 90 meters, while about 10% of flights terminate beyond 1 km. Although dispersal estimation from simulated epidemics suggests that these distances may be overestimated, this large number of flights terminating within some tens of meters of the source tree is consistent with previous studies of within-patch clustering of trees infected by PPV-M [32, 46, 47] or PPV-D [37, 48]. Indeed, in the latter study [37] 50% of the new PPV cases are shown to occur within 35-70 m of the nearest previous case; in addition, 10% of the new PPV cases were found beyond 200-460 m from the nearest previous case. Although the proportion of new PPV cases captured within a given radius is not equivalent to a dispersal kernel (e.g. because the trees are not always infected by the nearest previously detected neighbour), the figures are of the same order of magnitude. In particular, both studies highlight the long range of the dispersal kernel. Our estimation of the dispersal kernel at the landscape scale has important consequences. For example, the current French regulations enforce at least one visual inspection per year within 2.5 km of a detected sharka case (followed by the removal of all trees with sharka symptoms). Less than 3% of flights should thus go beyond this radius (Figure 6). In a patchy French landscape, most of these aphids would land outside a peach orchard and thus lead to no infection. Such procedures are thus likely to efficiently detect most of aphid-mediated secondary spread; actually, given the cost of surveillance and the speed of disease spread, this radius may even be oversized. Future work based on this study could aim at the definition of new management strategies against PPV. More generally, our results provide a unique reference point on the epidemiology, simulation and control of the principal group of plant viruses (i.e. those caused by nonpersistant aphid-borne viruses), which have a major epidemiological and economical impact. Finally, by focusing on incidence data the presented estimation approach is adaptable to many epidemiological situations, including other vector-borne and airborne fungal diseases.

## Acknowledgments

This work was supported by the EU (SharCo, FP7 204429), the Département Santé des Plantes et Environnement of INRA, and FranceAgriMer. The PPV-M dataset was provided by the Fédération Départementale de Défense Contre les Organismes Nuisibles. Sylain Grizard provided valuable technical assistance with Geographical Information Systems. We are also grateful to Annie Bouvier (INRA) for her technical support regarding the software CaliFloPP, and to Eric Montaudon and Véronique Martin (INRA) for their help with the MIGALE computer cluster.

